# Patient-derived tumor spheroid cultures as a promising tool to assist personalized therapeutic decisions in breast cancer

**DOI:** 10.1101/2021.06.26.450027

**Authors:** Sarah Hofmann, Raichel Cohen-Harazi, Yael Maizels, Igor Koman

**Affiliations:** Institute for Personalized and Translational Medicine, Ariel University, Ariel, Israel

**Keywords:** 3D models, personalized medicine, patient-derived spheroids, breast cancer

## Abstract

Breast cancer is the most common cause of cancer related death in women. Treatment of breast cancer has many limitations including a lack of accurate biomarkers to predict success of chemotherapy and intrinsic resistance of a significant group of patients to the gold standard of therapy. Therefore, new tools are needed to provide doctors with guidance in choosing the most effective treatment plan for a particular patient and thus to increase the survival rate for breast cancer patients.

Here, we present a successful method to grow in vitro spheroids from primary breast cancer tissue. Samples were received in accordance with relevant ethical guidelines and regulations. After tissue dissociation, in vitro spheroids were generated in a scaffold-free 96-well plate format. Spheroid composition was investigated by immunohistochemistry (IHC) of epithelial (Pan Cytokeratin (panCK)), stromal (Vimentin) and breast cancer-specific markers (ER, PR, HER2, GATA-3). Growth and cell viability of the spheroids were assessed upon treatment with multiple anti-cancer compounds. Student’s t-test and two-way ANOVA test were used to determine statistical significance.

We were able to successfully grow spheroids from 27 out of 31 samples from surgical resections of breast cancer tissue from previously untreated patients. Recapitulation of the histopathology of the tissue of origin was confirmed. Furthermore, a drug panel of standard first- and second-line chemotherapy drugs used to treat breast cancer was applied to assess the viability of the patient-derived spheroids and revealed variation between samples in the response of the spheroids to different drug treatments.

We investigated the feasibility and the utility of an in vitro patient-derived spheroid model for breast cancer therapy, and we conclude that spheroids serve as a highly effective platform to explore cancer therapeutics and personalized treatment efficacy. These results have significant implications for the application of this model in clinical personalized medicine.

## Introduction

Breast cancer is the most common cause of cancer related deaths for women worldwide. In Europe alone, approximately 630,000 women died from breast cancer in 2018 [1]. According to current European and American guidelines for treatment, breast cancers are categorized into molecular subtypes based on the expression of hormone receptors PR and ER as well as HER2 and Ki67. The subtypes include luminal A, luminal B, HER2+/non-luminal, and basal-like/triple negative, and each subtype has its own course of treatment with endocrine therapy recommended for hormone receptor positive patients and anti HER2 drugs for patients with high levels of HER2 expression [1, 2]. Yet, for the basal-like/triple negative subtype, where neither endocrine therapy or anti-HER2 therapy is appropriate, the first-line of therapy is classic chemotherapy which includes anthracyclines and/or taxanes [2–4].

While in breast cancer, there is a clear protocol for choosing a treatment plan for each patient, this treatment plan is, unfortunately, not always successful. Nearly 40% of women acquire tamoxifen resistance. Furthermore, intrinsically tamoxifen-resistant cancers often have a large number of gene alterations, making the choice of an alternative therapy anything but straightforward [5, 6]. In addition, there are no effective biomarkers to guide the choice of chemotherapy from a number of approved regimens. An unguided choice for first-line therapy can lead to a delay in effective treatment, and thus risk progression of the disease. Furthermore, each course of treatment is accompanied by suffering due to adverse side effects of chemotherapy [7, 8]. Even when actionable biomarkers do exist, they are not universally applicable and some patients who express a biomarker that is correlated with drug response do not respond to the predicted therapy. New tools to predict drug efficacy for individual patients would extend survival and prevent treatment with ineffective drugs.

One approach to developing new tools to guide individualized treatment selection utilizes chemotherapy sensitivity and resistance assays (CSRAs) which use viable tissue from a tumor to provide predictive information about response to treatment [9, 10]. A number of three dimensional (3D) models have been developed including tumor cells seeded in a matrix of extracellular proteins, multicellular tumor spheroids, organoids, tissue slices, bioreactors and microfluidic models [10–18]. The most efficient system to quickly and inexpensively culture patient-derived tissue in a 3D model are spheroids, scaffold-free, multicellular spheres containing cancer cells. These spheroids enable cell-cell interactions and can facilitate the production of the endogenous extracellular matrix to provide local tumor microenvironment-like conditions. Additional environmental considerations, including the access to nutrients, oxygen, growth factors, metabolites and paracrine factors, are also recapitulated [11, 18]. Unlike organoids, there is no need for expensive reagents such as specialized cell media factors or a basement membrane matrix (such as Matrigel) that can complicate downstream analysis of the tissue. Recently, it was shown that patient-derived spheroids could be grown from ovarian tumors and that these spheroids could accurately predict response to first-line therapy with 89% accuracy [19]. This clearly demonstrates the potential of this method to facilitate treatment choice, as well as to explore non-standard therapies, both of which would prevent a delay in effective treatment, eliminate unnecessary suffering, and ultimately improve prognosis.

To date, spheroids have been grown from a variety of breast cancer cell lines [20–24]. Yet, successfully growing spheroids from breast cancer patient samples has proven to be a challenge. This is due to a number of factors including the small size of breast tumors, the low number of viable cells which can be extracted and the heterogeneity of breast tissue which includes a high level of non-malignant tissue including adipose tissue [25, 26]. We have only seen two publications from a single research group where spheroids have been successfully grown from patient tumor tissue [26, 27].

Here, we present a successful method to grow in vitro spheroids from patient-derived breast tumor tissue. Our method is low cost, suited to drug screening, rapid and applicable to a large variety of breast cancer subtypes with nearly 90% success. This represents a feasible model for drug efficacy prediction for breast cancer patients that can help guide treatment decision making.

## Materials and methods

### Ethical approval and collection of patient samples

Ethical approval for the research was obtained from the Institutional Helsinki Committee at the Barzilai University Medical Center in Ashkelon, Israel according to the ICH-GCP guidelines (protocol No. 0282-18-TLV) under the 1964 Helsinki declaration and its later amendments or comparable ethical standards. Informed written consent to have data/samples from their medical records used in research, was obtained from all participants and/or their legal guardians and all research was performed in accordance with relevant guidelines and regulations. All tissue samples were collected between August 2018 until December 2019. Breast cancer samples were fully anonymized by the Institutional Biobank at Tel Aviv Sourasky Medical Center, Israel directly after surgery and were transported to our laboratory on ice in sterile tubes containing HypoThermosol® FRS medium (StemCell Technologies). Tissue was processed within 72 hours of surgery. Sample and clinical information about the patient were acquired anonymously, with identifying information encoded at the clinical site. Medical records for retrospective study from selected patients were received six months after collection.

### Generation of spheroids

The tissue was placed in a sterile 10 cm dish, photographed, and cut into 1-3 mm^3 pieces using a sterile razor blade. Representative pieces were saved for IHC analysis of the original tissue. The rest was further minced. Eventually, all remaining tissue was placed in a falcon tube containing Advanced DMEM supplemented with 1% Glutamax, 1% HEPES 1M, 1% Pen/Strep (all Gibco), 1x primocin (InvivoGen), 0.5 mg/ml collagenase (Sigma) and 0.2 mg/ml DNAse I (Sigma), and incubated on an orbital shaker at 220 rpm, at 37°C for up to 16 hrs. Where necessary, an additional digestion with 1x TrypLE (Gibco) was performed for 10 min at 37°C. Then, red blood cells were eliminated using the BD Pharm Lyse™ Lysing Buffer (BD Biosciences) according to the manufacturer’s instructions. Finally, the cell suspension was strained over 40 um Corning® cell strainers (Corning), and the single cell yield was assessed using the LUNA™ cell counter. Spheroids were generated using either the 3D InSight™ (GravityTRAP™ and GravityPLUS™) Hanging Drop System (InSphero AG) or the Corning® ULA spheroid microplates (Corning CLS4515) according to the manufacturer’s instructions. 1000 patient-derived cells were seeded in co-culture with HDFs (Sigma 106-05A) in a ratio of 1:1 or 1:3 per well in a 96-well plate format. Spheroids were grown in Mammary Epithelial Basal Medium (Lonza) supplemented with 2 mM L-Glutamine, 1% Penicillin-Streptomycin, 1% Fetal Bovine Serum (all Gibco), 5 ug/ml insulin (PromoCell), 0.5 ug/ml Hydrocortisone, 20 ng/ml EGF, 20 ng/ml human FGF10, 1 U/ml Heparin, 50 uM L-Ascorbic acid, 50 ng/ml Cholera Toxin, 35 ug/ml BPE (all Sigma), 1x B27 supplement (Life Technologies) and 20 ng/ml beta-Estradiol (Sigma). Media was changed every 2-3 days and spheroid growth was monitored by light microscopy using the Zeiss Axio Observer fluorescent microscope. The diameter “d” of spheroids was determined using the measurement feature of ImageJ (Version 1.52a). The size of each spheroid was calculated as: area= π * ((d/2))^2.

### Histology and IHC

Spheroids and original tissue pieces were collected into 1.5 ml Eppendorf tubes, washed in 1x PBS and fixed with 4% Paraformaldehyde (Bar Naor, Ltd.) for 15 min at RT. All further histology and IHC steps were performed by the staff of PathoLab (Rehovot, Israel). H&E staining was done using the Tissue Tek Prisma device under standard conditions. IHC staining for specific markers was done using the Ventana BenchMark Ultra System. The antibodies that were used and the respective dilutions are listed in S2 Table.

### Drug panel and CellTiter-Glo® 3D Cell Viability Assay

Breast cancer patient-derived spheroids were cultured in InSphero 3D InSight™ plates for 4 days. The following compounds were applied in 5 replicates per treatment at the respective final concentration: 100 nM Pac (T7191), 384 uM 5-FU (F6627), 1 µM Epi (E9406), 20 mM Met (PHR1084, all Sigma Aldrich). Redosing was performed after 3 days, and viability of the spheroids was determined after 7 days of treatment with the CellTiter-Glo® 3D Cell Viability Assay (Promega G9682) according to the manufacturer’s instructions. Luminescence readout was read in CLARIOstar® Plus plate reader (BMG LABTECH).

### Statistical analysis

The growth curve of spheroids was determined from the spheroid area. The data is presented as percentage of the size on day 1 of treatment. 5 replicates were measured. Student’s t-test and two-way ANOVA followed by Dunnett’s test were performed for statistical analysis. Graphs showing mean values with SEM were generated using GraphPad Prism (Version 9.0.0 (121)).

For the cell viability assay the mean of absolute luminescence from 5 replicates was determined. Student’s t-test and two-way ANOVA followed by Dunnett’s test were performed for statistical analysis. Graphs showing individual values, and mean values with SEM were generated using GraphPad Prism (Version 9.0.0 (121)).

### Availability of data and materials

All clinical data from all patient-derived samples are included in this published article as S1 Table.

All generated or analyzed data sets in this study are available from the corresponding author on reasonable request.

## Results

### Generation of spheroids from surgical samples of human breast cancer tissue

In order to establish a working protocol to grow in vitro spheroids from human patient material, tissue samples were received post-surgery from previously untreated human breast cancer patients (Fig 1). After dissociation of the tissue into a single cell suspension, spheroids were generated using a scaffold-free approach in the 96-well plate format from InSphero 3D InSight™ (GravityTRAP™ and GravityPLUS™) Hanging Drop System or the Corning® Ultra-Low Attachment (ULA) spheroid microplates. We observed that the reconstitution of the tumor microenvironment with stromal cells using normal human fibroblasts was fundamental to ensure the generation of multicellular aggregates in stable tumor spheroids. This approach was based on previous work with breast cancer cell lines which also used normal human dermal fibroblasts [20]. The best results were achieved with a tumor cell-to-fibroblast ratio of 1:3 (Fig 2A), but a 1:1 ratio also produced smooth, round and tightly packed spheroids (Fig 2B and 2C). The fibroblasts are recognized by the epithelial cells, and serve as a scaffold to facilitate the self-assembly of the tumor cells.

**Fig 1.**
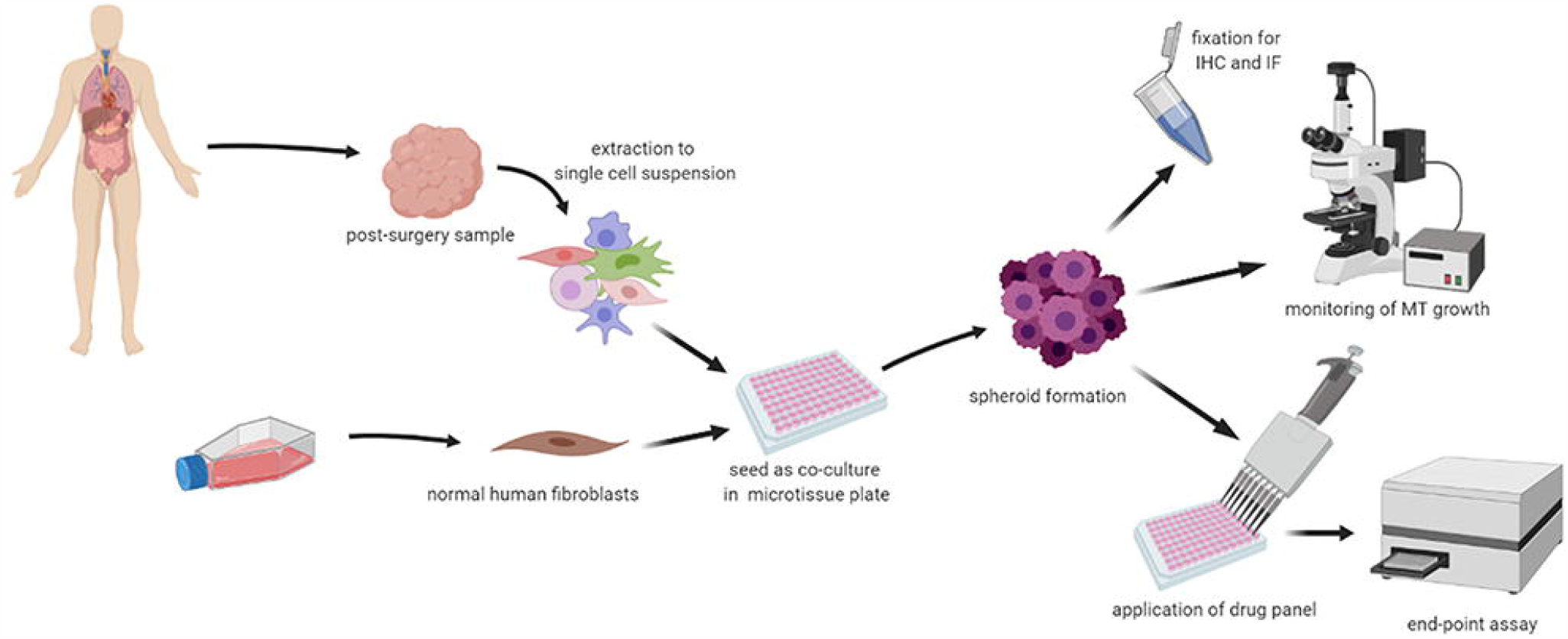
Workflow. Surgical resections of previously untreated tumors were dissociated into a single cell solution and seeded together with cultured normal human fibroblasts in specialized plates. Spheroids were maintained, monitored over time, and analyzed by a variety of assays including IHC assays or end-point cell viability measurements. (created with BioRender.com)

**Fig 2.**
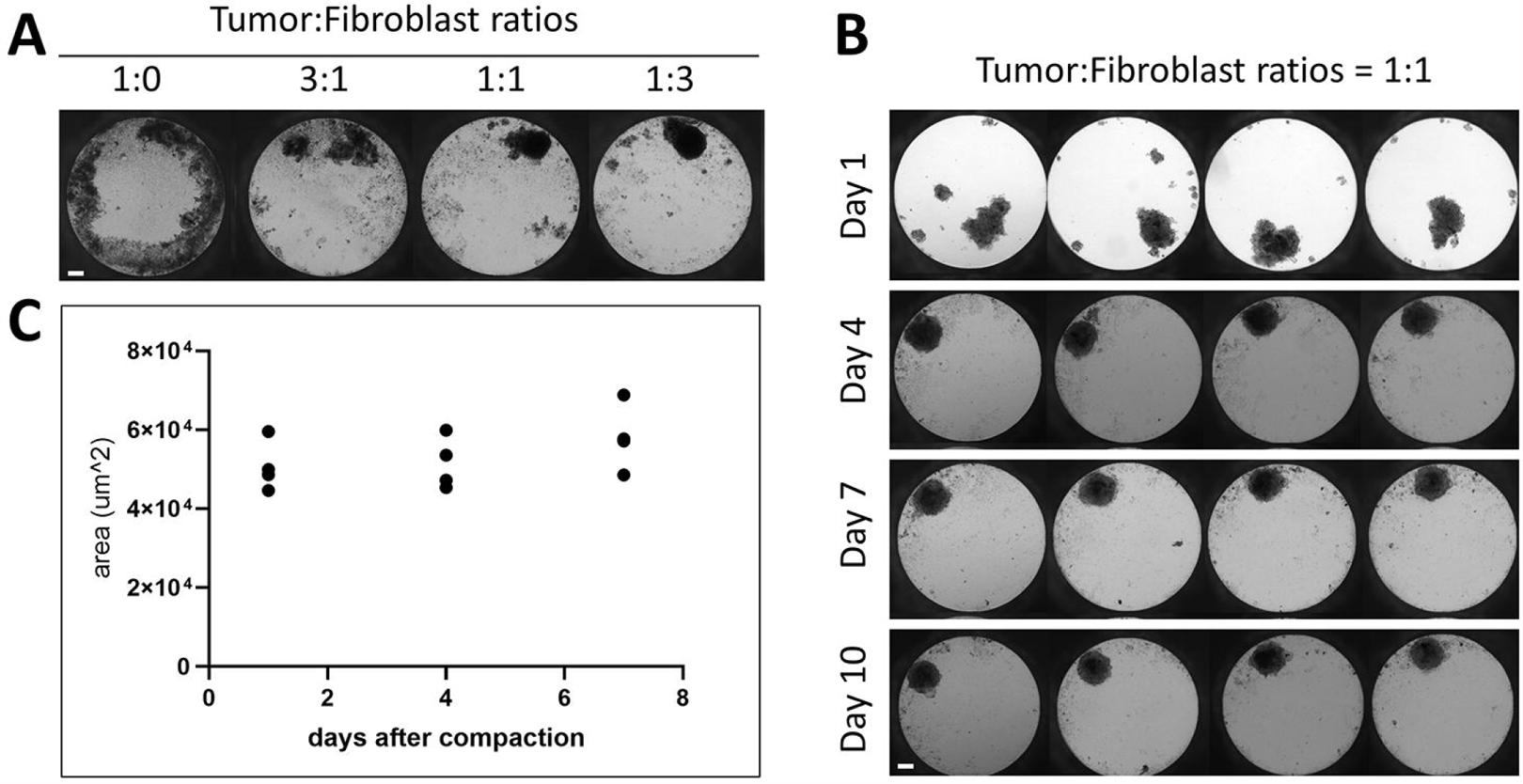
Under optimal growth conditions patient-derived tumor cells form tightly packed, round, uniform spheroids of similar size. (A) Tumor cells were seeded with varying ratios of normal human fibroblast. Spheroids were grown for seven days under optimal growth conditions. (B) Tumor cells were seeded 1:1 with normal fibroblasts, and growth was monitored over 10 days. (C) Spheroid size of four replicates grown over 7 days (after compaction at day 4) is shown. Size variations are similar between the replicates. Scale bar = 100 µm

On average, complete cell agglomeration into compact spheroids was seen after two to four days under optimal culture conditions (Fig 2B). The size variation between individual replicates of one patient sample is very low (Fig 2B and 2C). Once formed, spheroids could be fixed and stained for IHC or immunofluorescence (IF), monitored for growth by light microscopy and/or cell viability assays, or treated with a panel of chemotherapy drugs (Fig 1).

### Spheroids can be successfully grown across breast cancer tissues with distinct molecular signatures

We received surgical samples from a cohort of 31 breast cancer patients with varying genetic backgrounds and tumor grades. It is important to note that we were able to successfully grow spheroids from 27 out of the 31 samples, with an overall success rate of 87%. Our established criteria for success was the ability to grow at least 10 spheroids from the tumor tissue which would give us enough spheroids for control spheroids and drug testing of two different treatments. In 21 samples we were able to grow 20 spheroids and above which enabled us to test five different drugs or combinations. We had access to pathological and clinical data for most of the original tissue material. Table 1 shows a summary of the samples used in the study, with the available information regarding tumor stage and genetic background, as well as the success rate of establishing spheroids from the given tissues. We successfully grew spheroids from all genetic backgrounds and tumor grades received which included all possible genetic backgrounds for breast cancer with the exception of the HER2+/non-luminal type which we did not receive from the biobank. Spheroid growth success rate was independent of these tumor features based on the intrinsic data set. It is important to note that all samples from the triple-negative subtype were successfully grown using this procedure. This is encouraging since there is a lack of targeted drugs for this particular subtype [1, 28–30]. Thus, our established working protocol can generate in vitro spheroids from breast cancer tissues of different stages and genetic backgrounds.

**Table 1.** 3D spheroids were grown from breast cancer samples of varying stages and genetic backgrounds with an 87% success rate.

| **Tumor stage** | **I-II** | **II** | **II-III** | **III** | **N/A** | **Total** |
| --- | --- | --- | --- | --- | --- | --- |
| Total number of samples | 4 | 9 | 4 | 5 | 9 | 31 |
| Successful samples (%) | 4 (100%) | 7 (78%) | 4 (100%) | 4 (80%) | 8 (88%) | 27 (87%) |
| **Genetic background** | **Triple negative** | **Luminal A** | **Luminal B1** | **Luminal B2** | **N/A** | **Total** |
| Total number of samples | 4 | 11 | 9 | 5 | 2 | 31 |
| Successful samples (%) | 4 (100%) | 10 (91%) | 7 (78%) | 4 (80%) | 2 (100%) | 27 (87%) |
| **Ki67** | **0-10%** | **11-20%** | **21-40%** | **N/A** | **total** |  |
| Total number of samples | 17 | 6 | 3 | 5 | 31 |  |
| Successful samples (%) | 15 (88%) | 6 (100%) | 3 (100%) | 3 (60%) | 27 (87%) |  |

### The spheroids contain viable epithelial tumor tissue which maintains characteristics of the original tumor

The promising results achieved in generating in vitro spheroids from breast tumors of human origin, prompted us to determine the cellular composition of these spheroids. For this, spheroids from one patient cultured in the InSphero 3D InSight™ or Corning® ULA microplates, were fixed with paraformaldehyde and characterized by IHC. We found that spheroids generated from breast tissue contained both epithelial cells (as shown by panCK staining) and stromal cells (indicated by Vimentin staining) (Fig 3A and 3B). The epithelial cells in the spheroid are of tumor tissue origin, demonstrating that our technique supports the growth of original tumor tissue in the spheroid model. As expected, spheroids that were generated from human dermal fibroblasts (HDFs) alone only stained positive for Vimentin and not panCK (S1A and S1B Fig). It is important to note that we did not observe Vimentin-positive tumor cells. The tumor spheroids had clear Ki-67 staining indicating that the tumor-derived cells were actively proliferating in the spheroids (Fig 3C). In contrast, the HDF-only spheroids did not have Ki-67 staining (Fig 3C), showing that there is no concern that the fibroblasts proliferate and may become the bulk of the spheroid. Rather they mainly serve a structural and maintenance role. We show data from three representative spheroids from the same patient.

**Fig 3.**
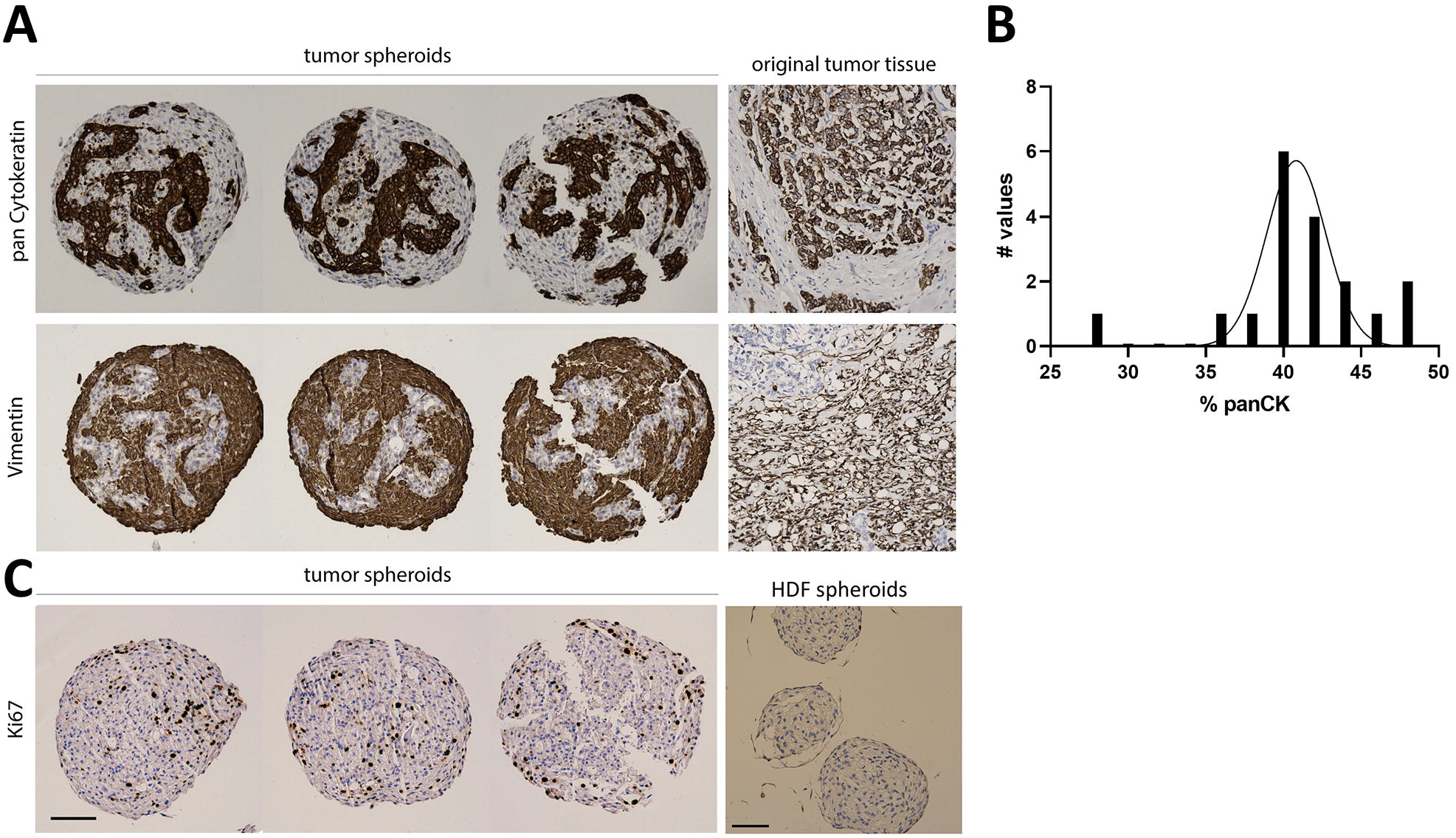
IHC staining of breast cancer spheroids reveals the distribution of fibroblasts and epithelial cells within the spheroids. (A) Three tumor spheroid replicates and the original tissue from one patient sample were stained by IHC for epithelial (panCK) and fibroblast (Vimentin). (B) Histogram analysis of panCK quantification shows that the majority of spheroids contains around 40-45% epithelial cells. (C) Three tumor spheroid replicates and HDF-only spheroids were stained by IHC for Ki-67. Positive Ki-67 staining is only observed in the tumor spheroids. Scale bar = 100 µm

Once we showed that viable tumor cells integrate into our spheroids, we investigated whether they maintain the molecular characteristics of the original tumor. For this investigation we used the markers most commonly used in the clinic to classify breast cancer and determine treatment, ER, PR and HER2 [1, 2]. Fig 4 shows a representative example of the same original tumor sample used in Fig 3, identified by clinical data as ER+, PR- and HER2- and classified as luminal B1 [1]. IHC staining of the spheroids from the same patient sample for these markers revealed that, like the original tissue, the spheroids were PR and HER2 negative (Fig 4). There was a significant number of ER+ cells in the original tumor. In the spheroids there was a small population of ER+ cells, this distribution is expected in light of the fact that the epithelial cells represent 25-50% of the spheroid content. After establishing that the spheroids reflected the staining pattern of the original tumor tissue, we examined another breast cancer marker, GATA-3, commonly expressed in over 90% of luminal B breast cancers [31]. GATA-3 was expressed in discrete cells in the original tumor and all three replicates of the tumor spheroids (Fig 4). Furthermore, we stained the HDF spheroids with the same panel of breast cancer markers and showed that they did not stain for any of the breast cancer markers (S1C Fig). This shows that expression of markers classically used to characterize breast cancer are maintained in the spheroid model and that this model recapitulates the original tissue in this respect.

**Fig 4.**
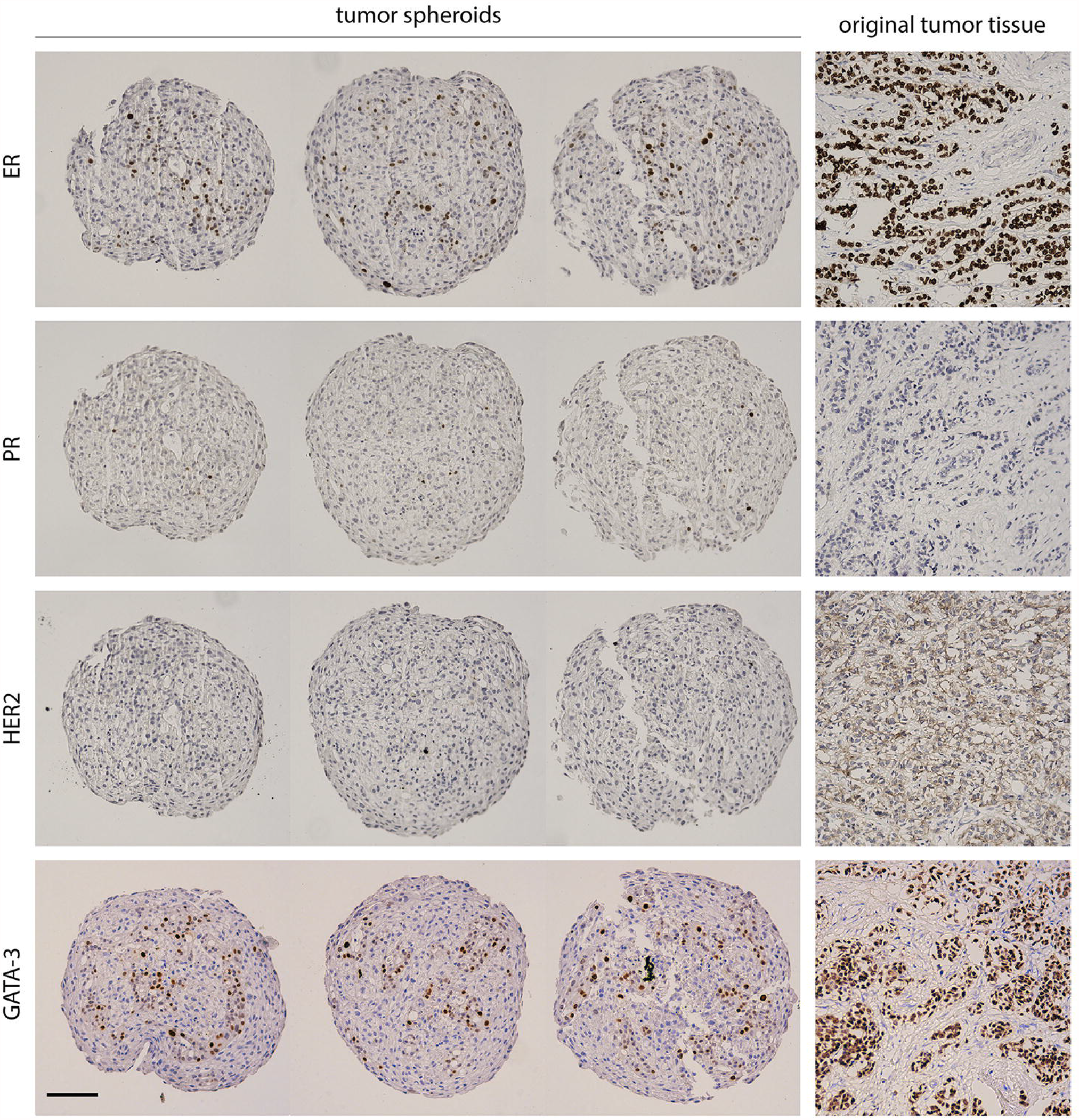
IHC staining of breast cancer spheroids reveals the heterogenous mixture of cellular components within the spheroids. Three tumor spheroid replicates and original tissue from one patient sample were stained by IHC for breast cancer markers (ER, PR, HER2 and GATA-3). The IHC staining of the spheroids resembles that of the original tissue (ER+, PR-, HER2-; luminal B1). Scale bar = 100 µm

### Spheroids respond to standard chemotherapy drugs

After successfully growing breast cancer spheroids from a variety of breast cancer patients and confirming that the spheroids reflect the molecular characteristics of the original tissue, we investigated whether this model could serve as a platform to test drug sensitivity. Therefore, to test the feasibility of the 3D spheroid model as a drug screening model, breast cancer patient-derived spheroids were treated with FDA-approved drugs. The list included Paclitaxel (Pac), Epirubicin (Epi), 5-Fluorouracil (5-FU) and Metformin (Met).

Spheroids were generated using the InSphero 3D InSight™ system. Following 4 days of incubation under optimal growth conditions, spheroids were treated with Pac, Epi, 5-FU, and Met for a period of up to one week (Fig 5A). The concentration of each drug was chosen from previous studies on 2D models and increased in accordance with the 3D setup [32–34]. Spheroid size was monitored by light microscopy, and the area was determined using ImageJ (Fig 5B). The size of the spheroids decreased significantly upon treatment with 5-FU (Fig 5C). Pac and Epi treatments also led to a visible reduction in spheroid size (Fig 5B). Following treatment, ATP levels were measured using a luminescence-based ATP assay (Fig 5D). As expected, ATP levels significantly decreased under treatment with the first- and second-line chemotherapy drugs for breast cancer, Pac, 5-FU and Epi, while Met showed no reduction in ATP levels (Fig 5D) or spheroid size (Fig 5B and 5C). Thus, the patient-derived spheroids respond, with expected variability, to clinically relevant chemotherapy drugs.

**Fig 5.**
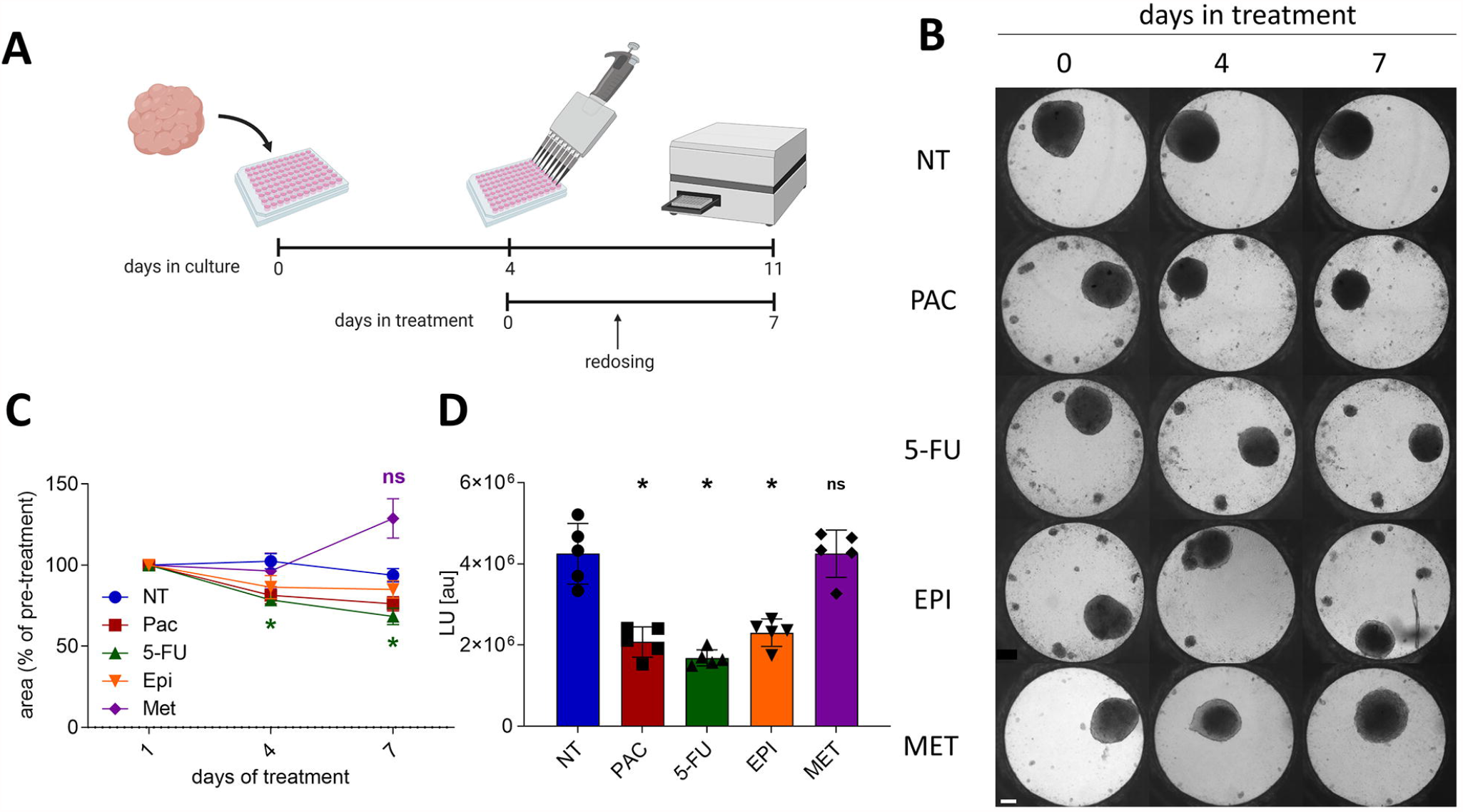
Drug panel reveals differences in response of breast cancer spheroids to multiple chemotherapeutic agents. (A) Schematic representation of experimental framework and timeline (created with BioRender.com). (B) Phase contrast images track morphological changes of spheroids generated from human breast cancer samples (ER+, PR+, HER2-, luminal A) upon a 7 day treatment in InSphero 3D InSight™ plates. Drug treatment included 100 nM Pac, 50 µg/ml 5-FU, 1 µM Epi or 20 mM Met and was applied 4 days after cell seeding, in 5 replicates per treatment. Redosing was performed after 3 days. Scale bar = 100 µm. (C) Size of tumor spheroids significantly decreases in response to 5-FU treatment. Data is presented as a percent of size on day 1 of treatment (n=5, asterisks indicate statistically significant differences from the NT; p<0.05, analyzed using two-way ANOVA, followed by Dunnett’s test). Mean values with SEM are shown. (D) Viability of the spheroids was determined after 7 days of treatment using a luminescence-based ATP assay. The average of absolute luminescence from 5 replicates is shown in the graph, with standard error. ATP levels significantly decreased under treatment with the chemotherapy drugs for breast cancer, Pac, 5-FU, and Epi, while Met showed no reduction in ATP levels (n=5, asterisks indicate statistically significant differences from the NT; p<0.0001, analyzed using two-way ANOVA, followed by Dunnett’s test). Mean values with SEM are shown.

## Discussion

### Improved method to generate spheroids from human breast cancer tissue

In this study we present an improved method for generating spheroids from cancerous human breast tissue. While spheroids have been derived from patient tissue in a number of histotypes their generation from breast tumor tissue has proved challenging [25, 26, 35, 36]. We are aware of only two publications where breast cancer spheroids have been successfully grown from patient samples [26, 37]. Here, we used material derived from a cohort of breast cancer patients to establish an effective protocol for the generation of spheroids utilizing a scaffold-free approach and tested their applicability in a variety of end-point assays. We employed a generic growth media that was supplemented with necessary growth factors and additives, such as insulin, heparin or hydrocortisone, as well as beta-estradiol. However, we refrained from adding stem cell niche factors, in order to avoid selective growth of only one particular subpopulation [38]. As described, stromal cells are indispensable to the 3D architecture of the tissue, contributing to the physical cell organization and to the biochemical signaling within the tumor microenvironment, and thus support key properties of solid tumors. Fostering cell-cell interactions between normal fibroblasts and cancer epithelial cells within the spheroids is therefore critically important for mimicking the tumor microenvironment in vitro [39–43]. On the other hand, excessive proliferation of the fibroblasts could threaten the overall composition of the spheroid. For this reason, we carefully selected the supplements of the growth media. For example, FGF10 has been reported to selectively promote epithelial cancer cell growth, while cholera toxin reduces the expansion of fibroblasts [15, 44, 45]. In addition, in most cases, the spheroids were cultured for a maximum of 2 weeks in medium with a low percentage of serum, thereby also limiting the expansion of the fibroblast population. Ki-67 staining of HDF and tumor spheroids demonstrated that the fibroblasts were not an actively replicating population in the spheroids (Fig 3C).

Our success rate in growing spheroids from primary tissue was at 87%, which is significantly higher than previously reported 73% and 59% for work on patient-derived spheroids in breast cancer [26, 27]. Spheroids which failed to grow (4 out of 31) were mostly due to contamination with bacteria or fungi (in 3 cases), or low quality of the original tissue piece which resulted in a low yield of viable epithelial cells after single cell extraction (in 1 case). This high success rate means that this system could be applicable for the vast majority of breast cancer patients.

### Validation of spheroid composition

In light of the heterogeneity of breast tumor tissue it was essential to ensure that the spheroids contained epithelial tissue. Therefore, we performed IHC analysis to identify and discriminate between the epithelial and stromal cells. This type of analysis was not performed in the two studies of spheroids from patient breast cancer tissue mentioned above [26, 27]. It has previously been shown that in spheroids generated from co-cultures of cancer cell lines and fibroblast lines the epithelial cells are primarily localized to the periphery of the spheroid, whereas the fibroblast cells make up the core [39, 46]. Surprisingly, and in contrast to what we and others have seen in spheroids from co-cultured cell lines, we did not detect a clear and distinct localization of epithelial and stromal cells in the spheroids generated from patient-derived material (see Fig 3 as a representative image of our consistent observation of this phenomenon). We also performed Ki-67 staining which showed that a portion of the epithelial cells were replicating while the fibroblast cells were not. In conclusion, our patient-derived breast cancer spheroids were composed of a mixture of fibroblasts and epithelial tumor tissue with no clear compartmentalization of the two populations.

Furthermore, we showed that the composition of the spheroids recapitulates the molecular features of the original tissue with regards to the classic breast cancer markers ER, PR and HER2 (Fig 4). Since these markers are standard in the clinic to characterize the tumor, it is important to note that their expression was maintained in the spheroid model [1, 2]. We further characterized the spheroids by looking at an additional common breast cancer marker, GATA-3, and saw that this marker is expressed both in the original tissue and in the tumor spheroids. This type of investigation was not performed in the previous studies on breast cancer spheroids. These data provide an important piece in the validation of our method.

### Responsiveness of spheroids to drug panel

To provide a preliminary proof-of-concept that spheroids could, indeed, be utilized as in vitro models for the prediction of drug response in personalized medicine, we applied a small panel of commonly used chemotherapy and adjuvant drugs, including Pac, Epi, 5-FU, and Met onto spheroids generated from patient-derived breast cancer tissue. While the tumor sample we used was of the Luminal A subtype which is most commonly treated by endocrine therapy, single agent chemotherapy is often used as a second line of therapy in these cases or if there is high tumor burden [2]. Given significant levels of tamoxifen resistance in ER+ cancer (up to 40%) and recurrence of up to 22% in 5 years it is important to arm doctors with tools to determine treatment in the case of resistance or recurrence [5, 6, 47]. In future experiments we plan on testing the first-line recommendations for the tumor type on the spheroids. Our system could also be used as a low risk assay to determine individual response to more experimental drugs. For example, Met has recently been reported to provide a promising adjuvant treatment for prostate and colorectal cancer and is currently being investigated as adjuvant therapy for breast cancer in an active phase III clinical trial [48]. With our drug panel we showed the potential of our system to show drug response for classic chemotherapy and investigational drugs.

As a readout for cell viability of the spheroids upon drug treatment, we chose a simple size assessment from bright-field images, as well as a luminescence-based ATP assay. We show that the reduction in spheroid size correlates with a decrease in intracellular ATP levels. All tested first-line chemotherapy drugs for breast cancer affected the viability of the spheroids, as expected. Similar drug panel assays in other contexts consist of dose-response data of compounds to estimate an appropriate dose for each drug regimen. Due to the relatively low yield of viable single cells extracted from breast cancer tissues, it is not possible at this time to generate enough spheroids to perform such analysis. However, in general, there are many challenges in the translation of doses determined in vitro to clinical dosing. The primary goal of a personalized medicine assay is foremost to assess the drugs and drug combinations to which an individual tumor is sensitive, rather than to provide exact dosing information.

Here, our main focus was to investigate spheroids for their variability and sensitivity to single agent application of specific drugs that were selected based on their clinical relevance for the tumor type. This valid, biological platform can significantly contribute to the choice of first-line treatment where single or combinatorial drugs are used for a better therapeutic response in individual patients. Shuford et al. have recently reported that spheroids generated from ovarian tissues accurately predicted the response and non-response of ovarian cancer patients to specific treatments.

### Summary: benefits and limitations of the spheroid model

3D models provide enormous benefit beyond conventional 2D cell culture models with regards to establishing an efficient in vitro pre-clinical platform for drug screening that accurately represents the original tumor [10, 19, 42]. Several 3D models have been developed including organoids, tissue slices, hydrogels, bioreactors, microfluidic models and scaffold-free spheroids [10–14, 16–18, 35]. In our research, we investigated the utility of spheroids as a model for drug testing on patient-derived breast cancer tissue. One of the primary limitations, that is true of all 3D culture methods, is the unpredictable quality of human biopsies and surgical resections as source of starting material. The tissue heterogeneity and size differ between various tissue types. Ultimately, breast tissue contains a high percentage of fat cells and a relatively low volume of epithelial cells, whereas other tissue types, such as colon or prostate yield a higher number of viable cancer cells [25]. Despite the challenges presented with the starting tissue in breast cancer we were able to generate spheroids with the relatively high success rate of nearly 90%. Before this model can be firmly established as a viable system for prediction of drug efficacy a more thorough investigation of the subpopulations that exist in the original tumor and the spheroids must be performed. Notably, the biggest advantage of our method, which utilizes the Corning® ULA microplates and the InSphero 3D InSight™ system is the straightforward seeding procedure and the ease of plate handling. In addition, within only a few days, spheroids are already visible and functional for drug treatment and respective end-point assays. Therefore, they provide a promising tool for pre-clinical prediction of drug response in personalized cancer therapy.

Overall, our success rate in generating spheroids from nearly 90% of the breast cancer tissue samples obtained, as well as the rapid time frame, use of minimal equipment and specialized reagents, and flexibility, support the potential of this method for clinical application. We were able to perform fifteen drug panels with at least one standard of care drug and we were consistently able to show a patient specific response. Our system, therefore, has the potential to be utilized as a tool for personalized prediction of the responsiveness of individual cancerous tissue to selected chemotherapeutic options. Further optimization and validation of our protocol will be performed in future pro- and retrospective studies.

## Acknowledgement

We thank Olga Shovman, Nili Turian, Karin Abeav, Hadas Fulman-Levy, and Tzachi Morad for technical assistance, and Drs. Sharona Even Ram and Gabriela Rozic for comments on the manuscript (all from the Institute for Personalized and Translational Medicine). We are grateful to Drs. Francesca Chiovaro, Wolfgang Moritz (both InSphero AG, Switzerland) and Dr. Sumeer Dhar for helpful discussions and comments on the study and the manuscript. We also thank Dr. Ayelet Izhaki and coworkers from the Institutional Biobank at Tel Aviv Sourasky Medical Center for help in collecting surgical specimens.

## Authors’ contributions

Conceptualization, RCH, SH and YM; methodology, RCH and SH; formal analysis, SH and YM; investigation, SH; writing—original draft preparation, RCH, SH; writing—review and editing, RCH, SH and YM; visualization, SH; supervision, IK; project administration, RCH, SH and YM; funding acquisition, IK. All authors read and approved the final manuscript.

## Supporting information

**S1 Fig. IHC staining reveals that HDF spheroids do not contain epithelial cells or express the breast cancer markers ER, PR, HER2 and GATA-3**.

(A) HDF spheroids were stained by IHC for epithelial marker (panCK) and fibroblast marker (Vimentin). (B) Histogram analysis of panCK quantification shows that the HDF spheroids do not contain epithelial cells. p value < 0.0001. (C) HDF spheroids were stained by IHC for the breast cancer markers ER, PR, HER2 and GATA-3. HDF spheroids show no positive staining for any of the breast cancer markers. Scale bar = 100 µm

**S1 Table. Clinical data of all patient-derived samples used in this study**.

**S2 Table. Antibodies used in this study for IHC**.

## Abbreviations

5-FU: 5-Fluorouracil
CSRAs: Chemotherapy sensitivity and resistance assays
Doxo: Doxorubicin
Epi: Epirubicin
HDF: Human dermal fibroblasts
IF: Immunofluorescence
IHC: Immunohistochemistry
Met: Metformin
Pac: Paclitaxel
panCK: Pan Cytokeratin
RTU: Ready to use
3D: Three dimensional
2D: Two dimensional
ULA: Ultra-Low Attachment

