## Supplementary figures and images for "Patient-derived tumor spheroid cultures as a promising tool to assist personalized therapeutic decisions in breast cancer"

### Supplement Figure S1

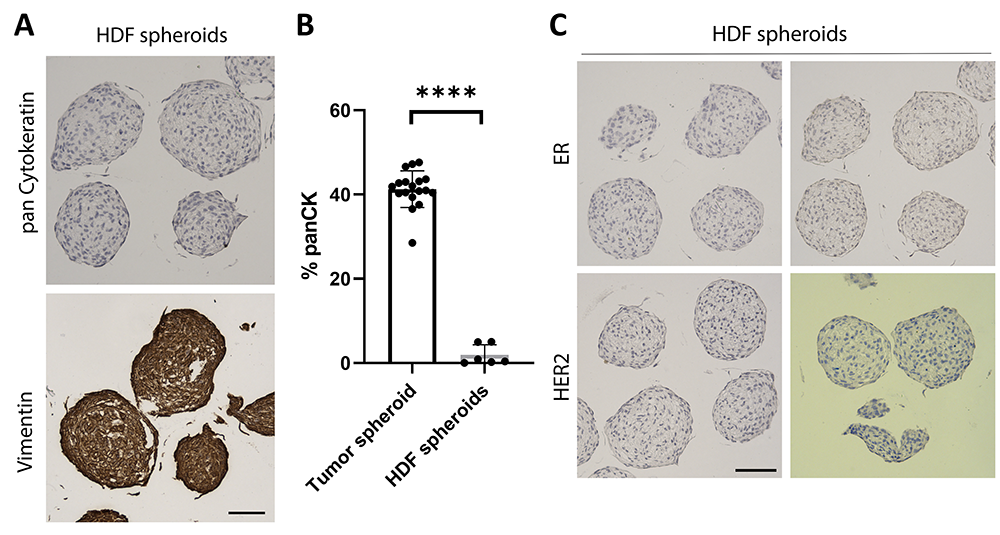
