## Supplement Table S1 for "Patient-derived tumor spheroid cultures as a promising tool to assist personalized therapeutic decisions in breast cancer"

### S1 Table. Clinical data of all patient-derived samples used in this study

| Patient | Tumor Grade | Successful growth of spheroids Yes (Y), No (N) | Genetic background | Subtype (T: triple neg; A: luminal A, Bi: luminal B(i); Bii: luminal B(ii) |
|---|---|---|---|---|
| 1 | 3 | Y | ER+ PR+ HER2+ Ki67 15% | Bii |
| 2 | N/A | Y | ER+ PR+ HER2- Ki67 8% | A |
| 3 | N/A | N | ER+ PR+ HER2+ Ki67 3% | Bii |
| 4 | 3 | Y | ER- PR- HER2- Ki67 40% | T |
| 5 | 2 | Y | ER+ PR+ HER2- Ki67 20% | A |
| 6 | 2 | Y | ER+ PR- HER2- Ki67 10% | Bi |
| 7 | 2 | Y | ER+ PR+ HER2- Ki67 5% | A |
| 8 | 1-2 | Y | ER+ PR+ HER2 N/A Ki67 N/A | N/A |
| 9 | 2 | Y | ER+ PR+ HER2- Ki67 4% | A |
| 10 | 2-3 | Y | ER+ PR- HER2- Ki67 N/A | Bi |
| 11 | N/A | Y | N/A | N/A |
| 12 | 2 | N | ER+ PR- HER2- Ki67 N/A | Bi |
| 13 | 1-2 | Y | ER+ PR- HER2- Ki67 3% | Bi |
| 14 | 2-3 | Y | ER+ PR+ HER2- Ki67 4% | A |
| 15 | N/A | Y | ER+ PR+ HER2- Ki67 5% | A |
| 16 | 2 | Y | ER+ PR+ HER2- Ki67 6% | A |
| 17 | 2 | Y | ER- PR- HER2- Ki67 10-15% | T |
| 18 | 2-3 | Y | ER+ PR- HER2- Ki67 2% | Bi |
| 19 | 1-2 | Y | ER+ PR+ HER2- Ki67 2% | A |
| 20 | N/A | Y | ER+ PR+ HER2+ Ki67 15% | Bii |
| 21 | N/A | Y | ER+ PR- HER2- Ki67 20% | Bi |
| 22 | 3 | Y | ER+ PR+ HER2+ Ki67 40% | Bii |
| 23 | 3 | N | ER+ PR- HER2- Ki67 N/A | Bi |
| 24 | 2-3 | Y | ER+ PR- HER2- Ki67 4% | Bi |
| 25 | NA | Y | ER+ PR+ HER2- Ki67 5-7% | A |
| 26 | 3 | Y | ER+ PR- HER2- Ki67 6-7% | Bi |
| 27 | 1-2 | Y | ER- PR- HER2- Ki67 10% | T |
| 28 | 2 | N | ER+ PR+ HER2- Ki67 5% | A |

| | | | | |
|---|---|---|---|---|
| 29 | N/A | Y | ER+ PR+ HER2- Ki67 7% | A |
| 30 | N/A | Y | ER- PR- HER2- Ki67 40% | T |
| 31 | 2 | Y | ER+ PR- HER2+ Ki67 20% | Bii |
