## Supplement Table S2 for "Patient-derived tumor spheroid cultures as a promising tool to assist personalized therapeutic decisions in breast cancer"

### S2 Table. Antibodies used in this study for IHC

| Marker | Antibody | Catalogue number | Dilution |
|---|---|---|---|
| ER | Rabbit monoclonal anti-Human ER, clone SP1 | Ventana cat# 790-4325 | RTU |
| PR | Mouse monoclonal anti-Human PRA, clone 16 | Leica Cat# NCL-L-PGR-312 | 1:100 |
| Her2 | Rabbit monoclonal anti-Human Her2/new, clone 4B5 | Ventana cat# 790-2991 | RTU |
| CK | Mouse monoclonal anti-Human cytokeratin, clone AE1/AE3 | Dako cat#M3515 | 1:200 |
| Gata3 | Mouse monoclonal anti-Human Gata3, clone L50-823 | Zytomed, cat#BMS054 | RTU |
| Vimentin | Mouse monoclonal anti-Human Vimentin, clone V9 | Dako, cat#M0725 | 1:1000 |
